# PhyloMagnet: Fast and accurate screening of short-read meta-omics data using gene-centric phylogenetics

**DOI:** 10.1101/688465

**Authors:** Max E. Schön, Laura Eme, Thijs J.G. Ettema

## Abstract

**Motivation:** Metagenomic and metatranscriptomic sequencing analyses have become increasingly popular tools for producing massive amounts of short-read data, often used for the reconstruction of draft genomes or the detection of (active) genes in microbial communities. Unfortunately, sequence assemblies of such datasets generally remain a computationally challenging task. Frequently, researchers are only interested in a specific group of organisms or genes; yet, the assembly of multiple datasets only to identify candidate sequences for a specific question is sometimes prohibitively slow, forcing researchers to select a subset of available datasets to address their question. Here we present PhyloMagnet, a workflow to screen meta-omics datasets for taxa and genes of interest using gene-centric assembly and phylogenetic placement of sequences.

**Results:** Using PhyloMagnet, we could identify up to 87% of the genera in an *in vitro* mock community with variable abundances, while the false positive predictions per single gene tree ranged from 0% to 23%. When applied to a group of metagenomes for which a set of MAGs have been published, we could detect the majority of the taxonomic labels that the MAGs had been annotated with. In a metatranscriptomic setting the phylogenetic placement of assembled contigs corresponds to that of transcripts obtained from transcriptome assembly.

**Availability:** PhyloMagnet is built using Nextflow, available at github.com/maxemil/PhyloMagnet and is developed and tested on Linux. It is released under the open source GNU GPL license and documentation is available at phylomagnet.readthedocs.io. Version 0.5 of PhyloMagnet was used for all benchmarks experiments.

## 1 Introduction

High-throughput DNA sequencing has revolutionized biology, opening up new fields of research and enabling new fundamental insights in the life sciences. During the past decades, several sequencing technologies have been developed, each differing significantly in sequence read length, quality and throughput (Mardis, 2017). Applications comprise DNA shotgun sequencing as well as RNA sequencing of complex microbial communities, termed metagenomics and metatranscriptomics, respectively (Mitchell *et al.*, 2018).

Large environmental sequencing initiatives like the Tara Oceans project (Sunagawa *et al.*, 2015) have provided researchers with enormous amounts of metagenome data. Using recently developed genome-resolved or genome-centric metagenomic approaches, draft genomes or MAGs (Metagenome assembled genomes) of uncultured taxa can be assembled for the first time from shotgun metagenomic sequencing data of microbial communities (Alneberg *et al.*, 2014; Eren *et al.*, 2015). In order to apply those tools, however, metagenome assembly needs to be performed, which is computationally demanding and introduces additional challenges compared to single genome assembly such as the uneven coverage of contigs (contiguous sequences) from different organisms or the presence of micro-diversity (Quince *et al.*, 2017). Together with the ever-growing sequencing capacity, it becomes increasingly demanding to identify which of the available datasets (publicly deposited or locally generated sequence datasets) actually contain sequence data of a given taxon or gene of interest.

Instead of assembling short reads into longer contigs, the taxonomic composition of a metagenomic or metatranscriptomic dataset can be assessed using microbiome profilers that classify reads directly. In general, these tools base their classification on the comparison of reads to reference sequences with a known taxonomy, and either work similar to the BLAST algorithm (e.g. Huson *et al.*, 2016; Truong *et al.* 2015) or use exact k-mer matches to such reference sequences to classify reads (e.g. Ounit *et al.*, 2015; Wood and Salzberg, 2014). Development in this area is continuing in order to increase analysis speed while reducing memory footprint. Currently, DIAMOND is one of the fastest local aligners that has a sensitivity comparable to BLAST (Buchfink *et al.*, 2015), and MetaCache is one of the fastest and most memory efficient k-mer based classifiers, using only a discriminatory subset of available k-mers (Müller *et al.*, 2017). All of these approaches, however, are based on sequence similarity, which can be incongruent with the true phylogenetic relationship of sequences (Smith and Pease, 2017).

Traditional phylogenetic tools on the other hand offer several robust evolutionary models for both nucleic and amino acids that theoretically allow for a more reliable taxonomic assignment of sequences, but are slow compared to similarity-based methods, usually prohibiting their application to large metagenome datasets. In addition, short reads generally do not provide enough phylogenetic signal, leading to artifactual inferences (Matsen *et al.*, 2010). Several tools have been developed to overcome these barriers by instead placing fragmentary sequences (particularly from amplicon sequencing data) onto a phylogenetic reference tree (Matsen *et al.*, 2010; Berger *et al.*, 2011; Barbera *et al.*, 2019).

Shotgun metagenomic or metatranscriptomic data is often analysed with a focus on gene rather than genome reconstruction, and is then usually called gene-centric. In this approach, the short reads or the assembled sequences are partitioned according to their affiliation to gene families. These methods can be used to determine which genes are present or actively transcribed in a sample, and can be combined with assemblers to reconstruct full-length sequences for a gene of interest. There exist several gene-centric targeted assemblers that perform de-novo reconstruction, e.g. via an overlap graph of candidate reads (Kucuk *et al.*, 2017; Pericard *et al.*, 2017; Steinegger *et al.*, 2018; Gruber-Vodicka *et al.*, 2019). While several of those only reconstruct the 16S rRNA gene or are limited to transcriptomic data, the MEGAN gene-centric assembler reconstructs contigs based on the alignment of reads to any reference protein sequence (Huson *et al.*, 2017).

A recently published tool, GraftM, uses the ideas of phylogenetic placement and gene-centric metagenomics to taxonomically classify sequences of genes within metagenomes (Boyd *et al.*, 2018). It is capable of placing either short-read sequences or pre-assembled metagenomic contigs onto a single reference tree at a time, but does not perform gene-centric assembly, which would increase phylogenetic signal of query sequences. Additionally, its reference trees can only be inferred using the extremely fast but less accurate maximum-likelihood-based tree inference program FastTree (Price *et al.*, 2010; Zhou *et al.*, 2018). Here, we present PhyloMagnet, an efficient workflow management system for parallel handling of both references and queries, gene-centric assembly, and robust phylogenetic inference, and show that it outperforms GraftM in terms of runtime and classification precision and sensitivity.

The goals of the work presented here were to:

i. Create a computational workflow that could determine the presence of taxa of interest in large short-read datasets based on gene-centric assembly and robust phylogenetic inference, especially with the objective of selecting good candidate datasets for metagenomic assembly and genome-resolved metagenomics;
ii. Create a workflow that uses state-of-the-art methods and is versatile and fast enough to accommodate a broad range of applications, while being modular in order to easily incorporate new approaches;
iii. Compare the workflow’s performance in terms of computational footprint and sensitivity/precision to GraftM, another recently published tool with a similar application.

## 2 Implementation

PhyloMagnet exploits the idea of gene-centric assembly (Huson *et al.*, 2017) to efficiently screen sequence datasets of short reads for target genes, and to taxonomically classify assembled gene sequences using phylogenetic placement. Below is a description of the analysis steps employed by the pipeline (see also Fig. 1), which requires the following inputs:

a. One or several query short-read sequence data files in FASTQ or FASTA format (potentially ‘raw’, untrimmed reads, see 2.3), corresponding to the metagenomic or transcriptomic dataset(s) to query (Fig. 1:1);
b. One or several homologous groups of reference proteins, each sequence annotated with its taxonomic affiliation (in the EggNOG format, containing ncbi’s taxonomy ID and a unique identifier, e.g. ‘70448.Q0P3H7’).

**Figure 1:**
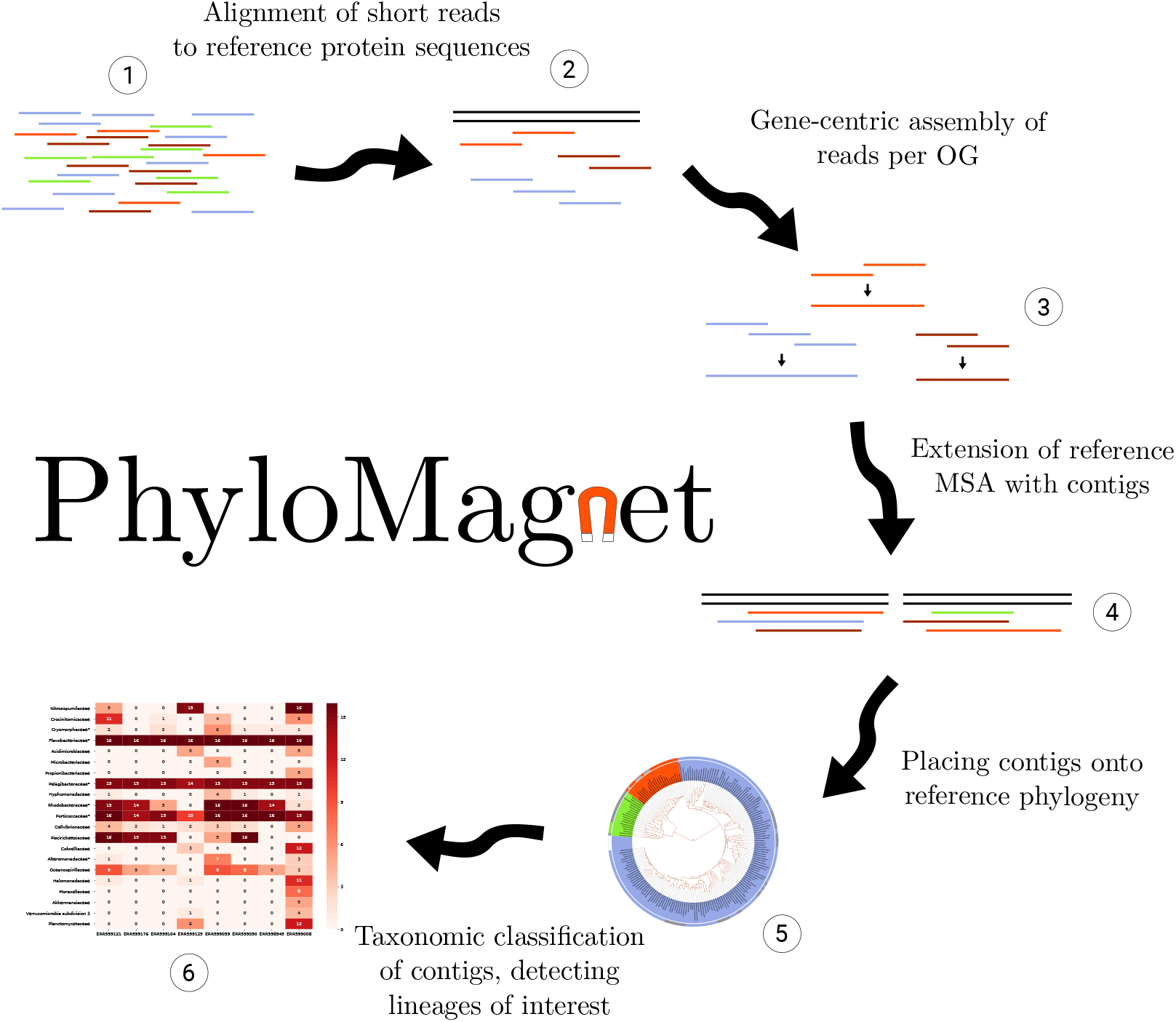
Illustration of the main steps in the PhyloMagnet workflow. (1) The required input is a dataset of short reads. (2) These reads are aligned against the complete set of protein references (blastX). (3) Using the protein alignments, homologous gene sequences are reconstructed for all groups of reference proteins. (4) The contigs are added to the reference protein alignments (5) and are subsequently placed onto the reference phylogenetic tree. (6) The results of the placement are summarized and the classification is visualized.

### 2.1 Alignment and tree reconstruction of reference

For each input group of reference sequences a multiple sequence alignment is computed using either MAFFT (Katoh and Standley, 2013) or PRANK (Löytynoja and Goldman, 2010), without applying any filtering or trimming methods. Then a reference tree is reconstructed using any of IQ-TREE (Nguyen *et al.*, 2015), RAxML (Stamatakis, 2014) or FastTree (Price *et al.*, 2010), making it possible to choose the appropriate method for a specific analysis. This way the user can make a trade-off between speed and quality of the reference tree and choose the appropriate evolutionary model. Reference alignments and trees can be precomputed (e.g. on a local machine) and then provided to PhyloMagnet as a compressed reference package (e.g. on an HPC cluster). This also increases reproducibility, as such reference packages can be released alongside results.

### 2.2 Alignment to reference protein sequences

Identifiers from the EggNOG database, containing orthologous groups of protein sequences from all domains of life with functional annotations (Huerta-Cepas *et al.*, 2016*b*), can be specified to be used as input reference sequences. Alternatively, sets of homologous protein sequences curated by the user in FASTA format can be used. In order to check for the potential presence of homologs encoding these proteins of interest in the query metagenomes or metatranscriptomes, each of the short-read datasets given as input (see 2.a above) is then aligned to the collection of reference protein sequences using the DIAMOND aligner in blastX mode (Fig. 1:2; Buchfink *et al.*, 2015).

### 2.3 Gene-centric assembly of reads

In a subsequent step, PhyloMagnet uses the gene-centric assembler implemented in MEGAN (Huson *et al.*, 2016, 2017) to assemble reads into contigs (Fig. 1:3). The assembly is performed independently for each orthologous group of reference proteins, and the available alignments of reads to the protein reference sequences of a group is used to infer overlaps between reads, thereby concatenating them into contigs. As only the aligned part (core) of each read is used for the assembly, no pre-processing such as adapter clipping or quality trimming is needed. The results are written to a FASTA file per orthologous group if any contig in that group passes the cut-off for the minimum length (200bp, can be adjusted if needed) that the gene-centric assembler uses. The assembled contigs are already in-frame and are subsequently translated into amino acid sequences using the standard genetic code.

### 2.4 Phylogenetic placement of reconstructed protein sequences

Next, the assembled and translated contigs are aligned to the alignments of each homologous reference group (maintaining the columns of the previously computed reference alignment), using the phylogeny-aware alignment tool PaPaRa (Fig. 1:4; Berger and Stamatakis, 2011). This alignment of reference sequences and contigs is then used to place the contigs onto the reference tree using the evolutionary placement algorithm (EPA-ng) (Fig. 1:5; Berger *et al.*, 2011; Barbera *et al.*, 2019). In a final stage the tool gappa is used to annotate the internal branches of the reference tree and assign taxonomic labels to the translated contigs based on the likelihood weights of the placement (Czech and Stamatakis, 2018). Then a summary list of taxonomic labels is created.

### 2.5 ‘Magnetizing’ trees and identifying candidate datasets

The user can choose to specify taxonomic names (e.g. ‘*Escherichia*’) that should be used to filter (‘magnetize’) the list of all labels, specify a taxonomic rank (e.g. ‘family’), or a combination of both. The occurrences of the chosen taxonomic labels are summarized per reference group and metagenomic or transcriptomic datasets in order to assist manual decision of candidate datasets (Fig. 1:6). The information of how many trees were positive for a taxon of interest can be used as an approximation of coverage (see 4). The user could for example select datasets that display differential coverage for subsequent genome extraction, which often relies on such differences to group genome contigs together (Albertsen *et al.*, 2013; Alneberg *et al.*, 2014).

### 2.6 Availability

PhyloMagnet is an open source software package and released under a GPLv3 license. It is written as a Nextflow (Di Tommaso *et al.*, 2017) script and available on github (github.com/maxemil/PhyloMagnet). Several functions and utilities are implemented either in python or bash (Dalke *et al.*, 2009; McKinney, 2010; Huerta-Cepas *et al.*, 2016*a*). All needed dependencies are available as a singularity (Kurtzer *et al.*, 2017) container (singularity-hub.org/collections/978) and the documentation can be found on ReadTheDocs (phylomagnet.readthedocs.io).

## 3 Benchmarking

To evaluate the performance of the PhyloMagnet workflow and exemplify its potential uses, we performed three benchmark experiments using an in vitro mock community as well as environmental metagenomic and metatranscriptomic sequencing datasets. We chose the datasets such that we could compare the results produced by PhyloMagnet to reference genome mapping data (Singer *et al.*, 2016), genomes extracted from metagenomes with taxonomic annotation (Delmont *et al.*, 2018) and an assembled metatranscriptome (Frazier *et al.*, 2017), respectively. For details on command line parameters see the supplementary methods.

### 3.1 Reference sequences

To assess the general taxonomic composition of datasets we used a set of 16 ribosomal proteins (rp16) that are thought to represent reliable phylogenetic markers, as they should be vertically inherited throughout evolution and present in a single copy in most organisms (Brown *et al.*, 2015). For this, we downloaded the corresponding sets of unaligned homologous sequences from the EggNOG database v4.5.1 (Huerta-Cepas *et al.*, 2016*b*).

As a second set of reference protein sequences, we used the set of 12 protein coding genes known to be present in chloroplast genomes of Dinophyceae (Howe *et al.*, 2008). This phylum of single-celled algae can be found in a wide range of aquatic environment and notably contains coral symbionts within the genus *Symbiodinium* (Gómez, 2012). For each of the genes we downloaded all available curated chloroplast encoded protein sequences for all phyla from UniProt (Apweiler *et al.*, 2004) as well as all available proteins from the Dinophyceae from the same database.

All reference groups were aligned using MAFFT E-INS-i (Katoh and Standley, 2013) and reference trees were reconstructed using IQ-TREE (under the LG+G+F model; Nguyen *et al.*, 2015).

### 3.2 Datasets

The first dataset we selected was the MBARC-26 (Mock Bacteria ARchaea Community), an in vitro mock community of 23 bacterial and 3 archaeal strains (in 24 genera) with finished reference genomes that were pooled and sequenced on an Illumina HiSeq instrument (Singer *et al.*, 2016). As the taxonomic classification is dependent on the reference sequences, we added orthologous sequences to the EggNOG rp16 references for those genera missing from the original EggNOG datasets. To avoid using identical sequences as references, we used available genomes from related species within the same genera to expand the rp16 references. The orthologous proteins were identified by performing HMMER (v3.1b2; Eddy, 2011) searches of the EggNOG rp16 reference alignments against the additional proteomes (see supplementary methods). These extended rp16 references were then used as references in PhyloMagnet in order to classify the MBARC-26 short-read data.

As a second dataset we used several metagenomic datasets from the geographic region ‘Southern Ocean’ that are part of the metagenomic datasets of the Tara Oceans Initiative, as defined by Delmont *et al.* (2018). We used the EggNOG rp16 references to assess taxonomic composition in those datasets and compared the results to the taxonomic classification of the MAGs reconstructed by Delmont *et al.* (2018). The authors extracted 375 genome bins from these datasets, but only presented detailed information, including taxonomy, for 13 ‘non-redundant’ MAGs that passed several quality and completeness filters. To be able to compare our classification results to a more extensive set of extracted genome bins, we inferred taxonomic labels for those bins that were not part of the 13 non-redundant MAGs using the tool sourmash that uses k-mer matches to taxonomically classify genomes (Titus Brown and Irber, 2016).

Finally, we analysed the metatranscriptomes published by Frazier *et al*. (2017), who sequenced mRNA from both healthy corals and such that are affected by so-called ‘bleaching’, a stress response in which *Symbiodinium* symbionts are expulsed (Howe *et al.*, 2008). We used the chloroplast protein references to search for Dinophyceae (and especially *Symbiodinium*) sequences in the metatranscriptome data. We then compared the assembled sequences and their placement in the reference trees with the sequences from the metatranscriptomic assembly available at NCBI’s GEO database (Barrett *et al.*, 2012). Similar to how the assembly was generated, we combined all of the 27 individual datasets by Frazier *et al*. (2017) into a single dataset for this analysis. In order to identify the transcripts in the assembly, we performed a tblastN search, querying the reference sequences against a database of the quality filtered transcripts. The identified sequences were then, analogous to how sequences are classified in PhyloMagnet, placed onto the reference tree with EPA-ng.

### 3.3 Comparison with GraftM

We compared the performance of PhyloMagnet with that of the recently published tool GraftM (Boyd *et al.*, 2018, v0.11.1). GraftM also places sequences (either unassembled reads or pre-assembled contigs) onto a reference phylogeny using the tool pplacer, for which EPA-ng represents a scalable replacement that is able to handle larger amounts of data. We created GraftM reference packages (gpkgs; containing the reference alignment, tree and the taxonomic annotation) from each of the extended rp16 references using the create command (see supplementary methods). We then used each gpkg to analyse the MBARC-26 dataset and recovered taxonomic classifications of the query sequences. For both tools, we counted the number of genera that were correctly identified in each tree (true positives) as well as the number of genera that were identified even though they were not present in the MBARC-26 mock community (false positives). We also assessed the runtime and memory consumption of both tools for analysis of the full MBARC-26 dataset (50 Gb) as well as for subsamples of 1% and 10% (0.5 and 5 Gb, respectively).

## 4 Results

### 4.1 Classification of ribosomal proteins in the MBARC-26 dataset

We evaluated the performance of PhyloMagnet and GraftM to detect the presence of the 24 MBARC genera (23 of those detectable, as Nocardiopsis was part of the pooled community but not present in the sequence data from Singer *et al.*, 2016) in the metagenomic dataset (Fig. S1 and Table S1). The number of correctly detected as well as falsely reported genera are displayed in Fig. 2. PhyloMagnet correctly identified up to 20 (87%) of the MBARC genera and up to 7 (with an average of 2) false positive genera in all of the 16 trees. In contrast, GraftM identified a maximum of 9 (39%) of the correct MBARC genera while giving up to 14 (with an average of 4) false positives for each tree (Fig. 2). Some of the reported false positive and false negative errors of both PhyloMagnet and GraftM could be attributed to closely related and possibly unresolved taxonomic groups such as *Escherichia*/*Salmonella*, *Thermobacillus*/*Paenibacillus* or possibly *Clostridium*/*Ruminiclostridium*. Another confounding factor might be the well-known disagreement between phylogeny and taxonomy in some cases (e.g. *Escherichia*/*Salmonella*; Retchless and Lawrence, 2010). Some taxa with very low abundance in the data (e.g. *Corynebacterium* and *Clostridium*) were picked up by GraftM but not PhyloMagnet, which is likely due to the fact that there are not enough reads to reconstruct longer contigs for theses taxa, impeding an identification by PhyloMagnet as we used the default cut-off implemented in the gene-centric assembler. In general we observe a correspondence between the percentage of mapped reads (Singer *et al.*, 2016) and the number of trees a genus was detected in (Fig. S2), suggesting that we can use the number of trees as a rough proxy for the abundance of a taxon in a dataset. When comparing results for the full dataset and the subsampled datasets, PhyloMagnet seems to profit immensely from the additional data, likely because the assembler can connect more reads and thus reconstruct more contigs above the length threshold. In terms of runtime, PhyloMagnet is twice as fast as GraftM on 10 CPUs, making more efficient use of available computational resources. It uses, however, significantly more memory due to the requirements of MEGAN that performs the memory intensive sequence assembly, which GraftM does not include (see Fig. S1).

**Figure 2:**
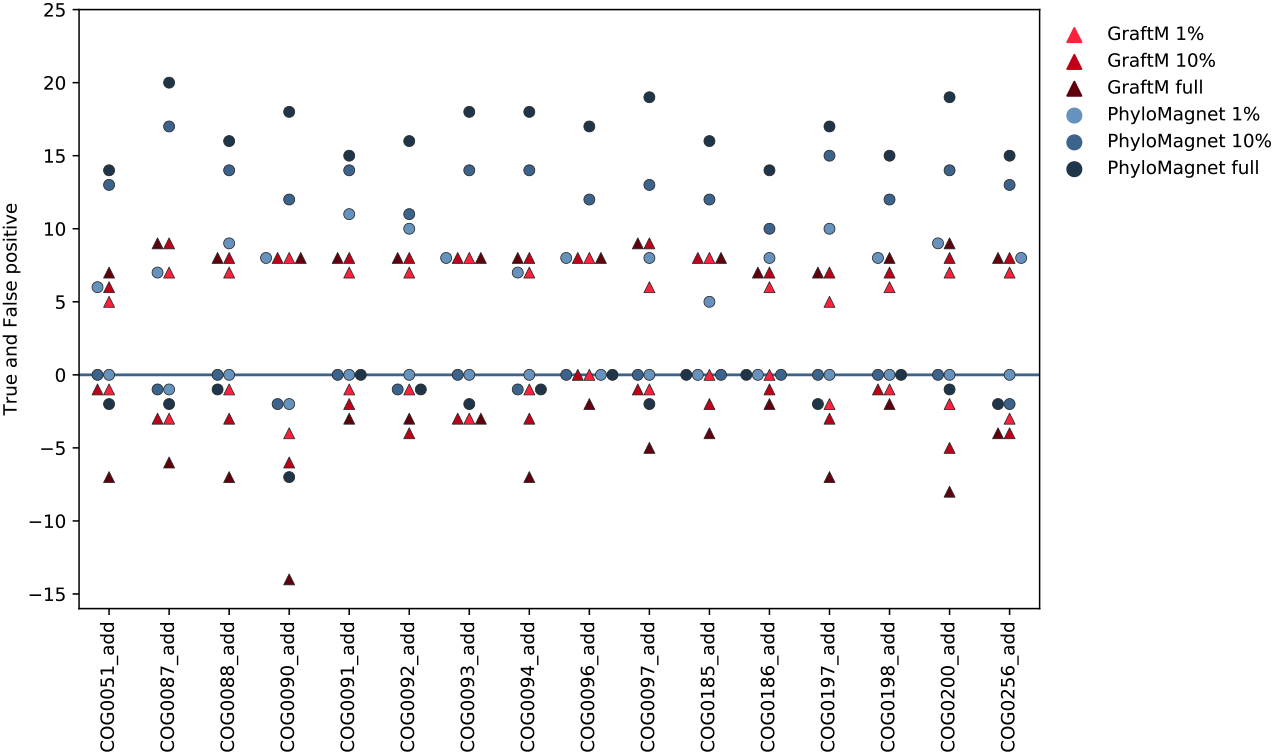
Classification results of PhyloMagnet and GraftM on the MBARC-26 dataset. True positive and false positive (on the negative y-axis including zero) values are shown for both PhyloMagnet (blue circles) and GraftM (red triangles) and for each reference OG (x-axis). The 3 different dataset sizes are shown by lighter (1%), middle (10%) and darker (full dataset) shades of the respective color.

### 4.2 Classification of taxa in the Tara Southern Ocean dataset

The PhyloMagnet workflow could identify 65 taxa (families) over the 8 datasets and 16 reference trees, whereof 21 were found in at least 4 trees for at least one dataset (Fig. 3). These taxa cover all but one of the taxonomic groups for which genomic bins could be identified by Delmont *et al*. (2018, marked with an asterisk in Fig. 3). Most noticeable, the authors of that study recovered 6 non-redundant high-quality MAGs for the family Flavobacteriaceae which could be identified in every single tree for each dataset here as well. Further, the original authors identified one MAG each within the Alteromonadaceae, the Rickettsiales and the Alphaproteobacteria as well as two within Gammaproteobacteria. All of these taxonomic groups were detected by PhyloMagnet except for the Rickettsiales, who could have been misidenfied as Pelagibacterales in the phylogenetic placement. Alternatively, the genomic bins could have been mislabelled as Rickettsiales and actually belong to the Pelagibacterales, as those two lineages commonly artifactually branch together in phylogenetic trees due to convergent genome streamlining resulting in a similar sequence composition bias (Roger *et al.*, 2017; Martijn *et al.*, 2018; Rodríguez-Ezpeleta and Embley, 2012; Viklund *et al.*, 2013). It is very likely that the MAGs that were labelled as Gammaproteobacteria by Delmont *et al*. (2018) are actually members of the Piscirickettsiaceae or Porticoccaceae, which were both detected by PhyloMagnet in several individual datasets and the majority of single gene trees. Here, we also recovered the additional taxonomic labels (Porticoccaceae, Rhodobacteraceae, Pelagibacteraceae, Cryomorphaceae) that could be assigned to raw genomic bins (which were not included in the original analyses as they did not pass quality and/or completeness thresholds) from the same study (Table S2). Some of the labels we recovered were not represented by any MAGs/Bins, indicating either false positive classification of contigs or a low abundance of the genomic DNA, such that no genome bins could be reconstructed by Delmont *et al*. (2018).

**Figure 3:**
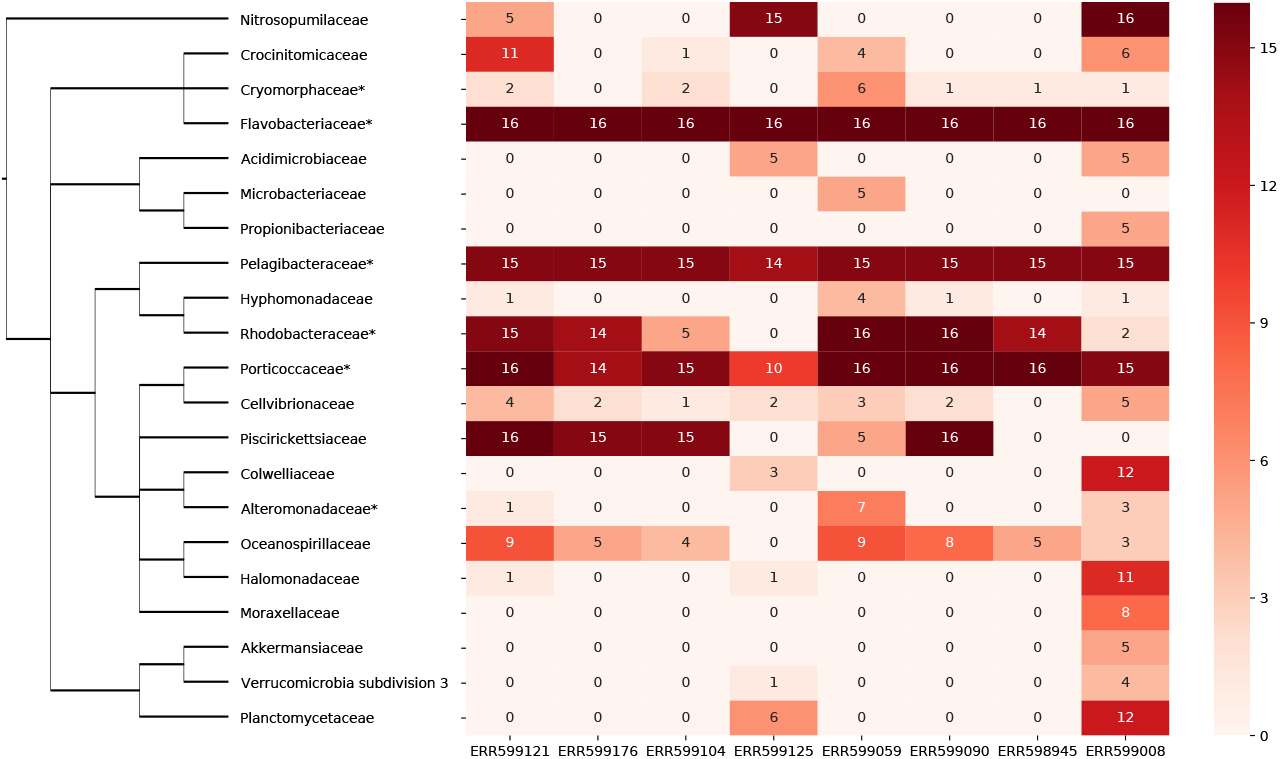
Classification results of the Tara Southern Oceans datasets. The heatmap on the right shows the identified taxa for each of the eight datasets, and only taxa which could be identified in at least four trees in at least one dataset are shown. For each combination of dataset and taxon, the value represents the number of trees a sequence from that dataset has been labelled with this taxon name. On the left a taxonomic tree (as defined by the ncbi taxonomy database) showing the relationship of lineages is depicted. Lineages represented by non-redundant and raw bins from Delmont *et al*. (2018) are marked with an asterisk.

### 4.3 Identification of chloroplast genes in the Coral bleaching dataset

Using PhyloMagnet contigs were reconstructed from the pooled coral bleaching dataset of Frazier *et al*. (2017). Using the Phylogenetic placement workflow, contigs classified as Symbiodiniaceae could be identified in 10 out of the 12 chloroplast gene reference trees. The number of contigs that were reconstructed for each gene from the pooled sequencing data of 23 datasets ranged from 2 (*psb*E) to as many as 169 (*psb*A), whereas we could identify either one or two transcripts from the corresponding published transcriptome assembly for 9 out of the 12 genes. The two genes for which no contigs could be reconstructed were *psb*I and *pet*D, both missing in the assembled transcriptome as well, which is likely due to two distinct issues. First, the *psb*I gene is only around 30 amino acid residues long, making contigs shorter than the default length cut-off of 200bp implemented in the gene-centric assembler. Besides, *psb*I has never been identified, experimentally or computationally, in any *Symbiodinium* species, but the identification within the Dinophyceae comes from the species *Amphidinium operculatum* (Nisbet *et al.*, 2004; Barbrook *et al.*, 2014). Second, it seems that the transcription level of *pet*D is quite low, so that very few reads would have been sequenced, making assembly of contigs or transcripts virtually impossible (Nisbet *et al.*, 2008). In those cases where transcripts could be identified, they were generally placed on the same branches or very close within the reference tree as were all of the corresponding contigs (Fig. 4).

**Figure 4:**
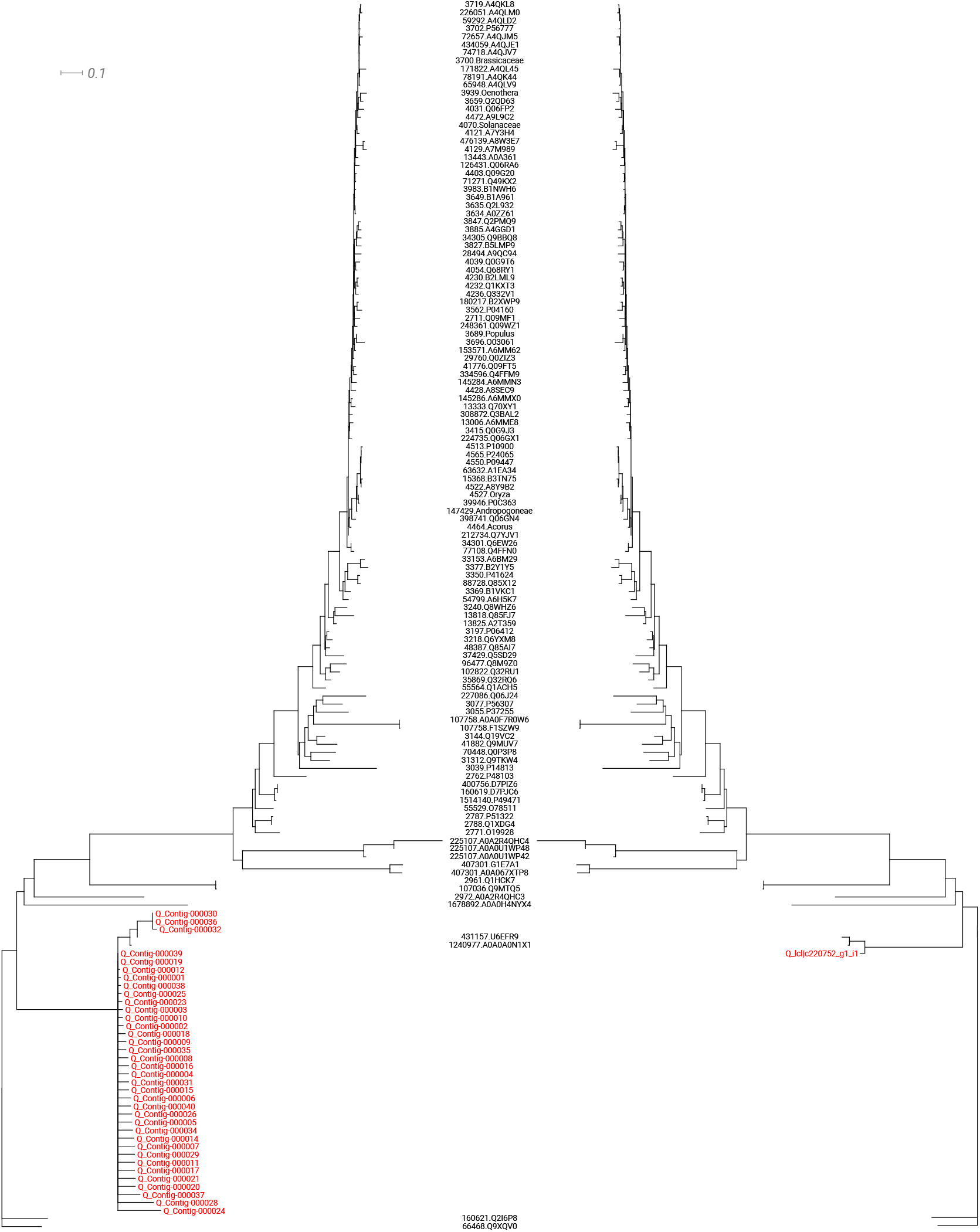
Phylogenetic placement of contigs and transcripts. As an example of the coral bleaching metatranscriptome analyses, the tree of the plastid gene *psb*B is shown. The sequences reconstructed by PhyloMagnet (left) and the assembled and quality-filtered transcripts from Frazier *et al*. (2017, right) were placed onto the reference tree. All sequences were placed on branches sister to two *Symbiodinium* sequences (U6EFR9 and A0A0A0N1X1).

## 5 Conclusion

We have shown that by applying phylogenetic placement methods to protein sequences that were reconstructed from short-read sequencing data, our PhyloMagnet workflow can accurately identify short-read sequence datasets that contain sequences for genes and taxa of interest. We compared PhyloMagnet to a similar tool that does not rely on using a gene-centric assembly approach and demonstrated that PhyloMagnet is faster and has a higher precision and sensitivity (at the price of consuming more memory). PhyloMagnet allows researchers to explore the microbial diversity of a specific clade, or to specifically assess the presence of a metabolic pathway of interest. For example, PhyloMagnet was able to identify several lineages from single gene trees that match the results of a genome-resolved metagenomic study, showcasing how our tool could be used to screen the contents of a metagenomic dataset before applying metagenome assembly and binning methods.

Finally, we have also shown that the gene-centric phylogenetic approach of PhyloMagnet can be successfully used to efficiently detect expressed genes of taxa of interest in metatranscriptomic datasets.

Hence, PhyloMagnet represents a powerful tool that will enable researchers to pre-screen large metagenomic and metatranscriptomics datasets prior to engaging in time and resource consuming computational analyses in their research.

## Acknowledgements

Several computations were performed on resources provided by SNIC through Uppsala Multidisciplinary Center for Advanced Computational Science (UPPMAX) under Project SNIC 2019-3-28.

## References

Albertsen,M., Hugenholtz,P., Skarshewski,A., Nielsen,K.L., Tyson,G.W. and Nielsen,P.H. (2013) Genome sequences of rare, uncultured bacteria obtained by differential coverage binning of multiple metagenomes. Nature Biotechnology, 31(6), 533–538.

Alneberg,J., Bjarnason,B.S., De Bruijn,I., Schirmer,M., Quick,J., Ijaz,U.Z., Lahti,L., Loman,N.J., Andersson,A.F. and Quince,C. (2014) Binning metagenomic contigs by coverage and composition. Nature Methods, 11(11), 1144–1146.

Apweiler,R., Bairoch,A., Wu,C.H., Barker,W.C., Boeckmann,B., Ferro,S., Gasteiger,E., Huang,H., Lopez,R., Magrane,M., Martin,M.J., Natale,D.A., O’Donovan,C., Redaschi,N. and Yeh,L.S.L. (2004) UniProt: the Universal Protein knowledgebase. Nucleic Acids Research, 32(90001), 115D–119.

Barbera,P., Kozlov,A.M., Czech,L., Morel,B., Darriba,D., Flouri,T. and Stamatakis,A. (2019) EPA-ng: Massively Parallel Evolutionary Placement of Genetic Sequences. Systematic Biology, 68(2), 365–369.

Barbrook,A.C., Voolstra,C.R. and Howe,C.J. (2014) The Chloroplast Genome of a Symbiodinium sp. Clade C3 Isolate. Protist, 165(1), 1–13.

Barrett,T., Wilhite,S.E., Ledoux,P., Evangelista,C., Kim,I.F., Tomashevsky,M., Marshall,K.A., Phillippy,K.H., Sherman,P.M., Holko,M., Yefanov,A., Lee,H., Zhang,N., Robertson,C.L., Serova,N., Davis,S. and Soboleva,A. (2012) NCBI GEO: archive for functional genomics data sets—update. Nucleic Acids Research, 41(D1), D991–D995.

Berger,S.A., Krompass,D. and Stamatakis,A. (2011) Performance, Accuracy, and Web Server for Evolutionary Placement of Short Sequence Reads under Maximum Likelihood. Syst. Biol, 60(3), 291–302.

Berger,S.A. and Stamatakis,A. (2011) Aligning short reads to reference alignments and trees. Bioinformatics, 27(15), 2068–2075.

Boyd,J.A., Woodcroft,B.J. and Tyson,G.W. (2018) GraftM: a tool for scalable, phylogenetically informed classification of genes within metagenomes. Nucleic Acids Research, 46(10), e59–e59.

Brown,C.T., Hug,L.A., Thomas,B.C., Sharon,I., Castelle,C.J., Singh,A., Wilkins,M.J., Wrighton,K.C., Williams,K.H. and Banfield,J.F. (2015) Unusual biology across a group comprising more than 15% of domain Bacteria. Nature, 523(7559), 208–211.

Buchfink,B., Xie,C. and Huson,D.H. (2015) Fast and sensitive protein alignment using DIAMOND. Nature methods, 12(1), 59–60.

Czech,L. and Stamatakis,A. (2018) Scalable methods for post-processing, visualizing, and analyzing phylogenetic placements. bioRxiv, , 1–36.

Dalke,A., Wilczynski,B., Chapman,B.A., Cox,C.J., Kauff,F., Friedberg,I., Chang,J.T., de Hoon,M.J.L., Cock,P.J.A., Hamelryck,T. and Antao,T. (2009) Biopython: freely available Python tools for computational molecular biology and bioinformatics. Bioinformatics, 25(11), 1422–1423.

Delmont,T.O., Quince,C., Shaiber,A., Esen,Ö.C., Lee,S.T., Rappé,M.S., MacLellan,S.L., Lücker,S. and Eren,A.M. (2018) Nitrogen-fixing populations of Planctomycetes and Proteobacteria are abundant in surface ocean metagenomes. Nature Microbiology, 3(7), 804–813.

Di Tommaso,P., Chatzou,M., Floden,E.W., Barja,P.P., Palumbo,E. and Notredame,C. (2017) Nextflow enables reproducible computational workflows. Nature Biotechnology, 35(4), 316–319.

Eddy,S.R. (2011) Accelerated Profile HMM Searches. PLoS Computational Biology, 7(10), e1002195.

Eren,A.M., Esen,Ö.C., Quince,C., Vineis,J.H., Morrison,H.G., Sogin,M.L. and Delmont,T.O. (2015) Anvi’o: an advanced analysis and visualization platform for ‘omics data. PeerJ, 3, e1319.

Frazier,M., Helmkampf,M., Bellinger,M.R., Geib,S.M. and Takabayashi,M. (2017) De novo metatranscriptome assembly and coral gene expression profile of Montipora capitata with growth anomaly. BMC Genomics, 18(1), 1–11.

Gómez,F. (2012) A quantitative review of the lifestyle, habitat and trophic diversity of dinoflagellates (Dinoflagellata, Alveolata). Systematics and Biodiversity, 10(3), 267–275.

Gruber-Vodicka,H.R., Seah,B.K. and Pruesse,E. (2019) phyloFlash — Rapid SSU rRNA profiling and targeted assembly from metagenomes. bioRxiv, , 521922.

Howe,C.J., Nisbet,R.E.R. and Barbrook,A.C. (2008) The remarkable chloroplast genome of dinoflagellates. Journal of Experimental Botany, 59(5), 1035–1045.

Huerta-Cepas,J., Serra,F. and Bork,P. (2016a) ETE 3: reconstruction, analysis, and visualization of phylogenomic data. Molecular Biology and Evolution, 33(6), 1635–1638.

Huerta-Cepas,J., Szklarczyk,D., Forslund,K., Cook,H., Heller,D., Walter,M.C., Rattei,T., Mende,D.R., Sunagawa,S., Kuhn,M., Jensen,L.J., Von Mering,C. and Bork,P. (2016b) EGGNOG 4.5: A hierarchical orthology framework with improved functional annotations for eukaryotic, prokaryotic and viral sequences. Nucleic Acids Research, 44(D1), D286–D293.

Huson,D.H., Beier,S., Flade,I., Górska,A., El-Hadidi,M., Mitra,S., Ruscheweyh,H.J. and Tappu,R. (2016) MEGAN community edition - interactive exploration and analysis of large-scale microbiome sequencing data. PLoS Computational Biology, 12(6), e1004957.

Huson,D.H., Tappu,R., Bazinet,A.L., Xie,C., Cummings,M.P., Nieselt,K. and Williams,R. (2017) Fast and simple protein-alignment-guided assembly of orthologous gene families from microbiome sequencing reads. Microbiome, 5(1), 11.

Katoh,K. and Standley,D.M. (2013) MAFFT multiple sequence alignment software version 7: Improvements in performance and usability. Molecular Biology and Evolution, 30(4), 772–780.

Kucuk,E., Chu,J., Vandervalk,B.P., Austin Hammond,S., Warren,R.L. and Birol,I. (2017) Kollector: transcript-informed, targeted de novo assembly of gene loci. Bioinformatics, 33(17), 2789–2789.

Kurtzer,G.M., Sochat,V. and Bauer,M.W. (2017) Singularity: Scientific containers for mobility of compute. PLoS ONE, 12(5).

Löytynoja,A. and Goldman,N. (2010) webPRANK: a phylogeny-aware multiple sequence aligner with interactive alignment browser. BMC bioinformatics, 11(1), 579.

Mardis,E.R. (2017) DNA sequencing technologies: 2006-2016. Nature Protocols, 12(2), 213–218.

Martijn,J., Vosseberg,J., Guy,L., Offre,P. and Ettema,T.J. (2018) Deep mitochondrial origin outside the sampled alphaproteobacteria. Nature, 557(7703), 101–105.

Matsen,F.a., Kodner,R.B. and Armbrust,E.V. (2010) pplacer: linear time maximum-likelihood and Bayesian phylogenetic placement of sequences onto a fixed reference tree. BMC bioinformatics, 11(1), 538.

McKinney,W. (2010) Data structures for statistical computing in python. In Proceedings of the 9th Python in Science Conference, (van der Walt,S. and Millman,J., eds), pp. 51 – 56.

Mitchell,A.L., Scheremetjew,M., Denise,H., Potter,S., Tarkowska,A., Qureshi,M., Salazar,G.A., Pesseat,S., Boland,M.A., Hunter,F.M., Ten Hoopen,P., Alako,B., Amid,C., Wilkinson,D.J., Curtis,T.P., Cochrane,G. and Finn,R.D. (2018) EBI Metagenomics in 2017: Enriching the analysis of microbial communities, from sequence reads to assemblies. Nucleic Acids Research, 46(D1), D726–D735.

Müller,A., Hundt,C., Hildebrandt,A., Hankeln,T. and Schmidt,B. (2017) MetaCache: context-aware classification of metagenomic reads using minhashing. Bioinformatics, 33(23), 3740–3748.

Nguyen,L.T., Schmidt,H.A., Von Haeseler,A. and Minh,B.Q. (2015) IQ-TREE: A fast and effective stochastic algorithm for estimating maximum-likelihood phylogenies. Molecular Biology and Evolution, 32(1), 268–274.

Nisbet,R.E.R., Hiller,R.G., Barry,E.R., Skene,P., Barbrook,A.C. and Howe,C.J. (2008) Transcript Analysis of Dinoflagellate Plastid Gene Minicircles. Protist, 159(1), 31–39.

Nisbet,R.E.R., Koumandou,V.L., Barbrook,A.C. and Howe,C.J. (2004) Novel plastid gene minicircles in the dinoflagellate Amphidinium operculatum. Gene, 331(1-2), 141–147.

Ounit,R., Wanamaker,S., Close,T.J. and Lonardi,S. (2015) CLARK: fast and accurate classification of metagenomic and genomic sequences using discriminative k-mers. BMC Genomics, 16(1), 236.

Pericard,P., Dufresne,Y., Couderc,L., Blanquart,S. and Touzet,H. (2017) MATAM: reconstruction of phylogenetic marker genes from short sequencing reads in metagenomes. Bioinformatics, 34(October 2017), 585–591.

Price,M.N., Dehal,P.S. and Arkin,A.P. (2010) FastTree 2 - Approximately maximum-likelihood trees for large alignments. PLoS ONE, 5(3), e9490.

Quince,C., Walker,A.W., Simpson,J.T., Loman,N.J. and Segata,N. (2017) Shotgun metagenomics, from sampling to analysis. Nature Biotechnology, 35(9), 833–844.

Retchless,A.C. and Lawrence,J.G. (2010) Phylogenetic incongruence arising from fragmented speciation in enteric bacteria. Proceedings of the National Academy of Sciences, 107(25), 11453–11458.

Rodríguez-Ezpeleta,N. and Embley,T.M. (2012) The SAR11 group of alpha-proteobacteria is not related to the origin of mitochondria. PloS one, 7(1), e30520.

Roger,A.J., Muñoz-Gómez,S.A. and Kamikawa,R. (2017) The Origin and Diversification of Mitochondria. Current Biology, 27(21), R1177–R1192.

Singer,E., Andreopoulos,B., Bowers,R.M., Lee,J., Deshpande,S., Chiniquy,J., Ciobanu,D., Klenk,H.P., Zane,M., Daum,C., Clum,A., Cheng,J.F., Copeland,A. and Woyke,T. (2016) Next generation sequencing data of a defined microbial mock community. Scientific Data, 3, 160081.

Smith,S.A. and Pease,J.B. (2017) Heterogeneous molecular processes among the causes of how sequence similarity scores can fail to recapitulate phylogeny. Briefings in Bioinformatics, 18(3), 451–457.

Stamatakis,A. (2014) RAxML version 8: A tool for phylogenetic analysis and post-analysis of large phylogenies. Bioinformatics, 30(9), 1312–1313.

Steinegger,M., Mirdita,M. and Soding,J. (2018) Protein-level assembly increases protein sequence recovery from metagenomic samples manyfold. bioRxiv, , 386110.

Sunagawa,S., Coelho,L.P., Chaffron,S., Kultima,J.R., Labadie,K., Salazar,G., Djahanschiri,B., Zeller,G., Mende,D.R., Alberti,A., Cornejo-Castillo,F.M., Costea,P.I., Cruaud,C., D’Ovidio,F., Engelen,S., Ferrera,I., Gasol,J.M., Guidi,L., Hildebrand,F., Kokoszka,F., Lepoivre,C., Lima-Mendez,G., Poulain,J., Poulos,B.T., Royo-Llonch,M., Sarmento,H., Vieira-Silva,S., Dimier,C., Picheral,M., Searson,S., Kandels-Lewis,S., Bowler,C., de Vargas,C., Gorsky,G., Grimsley,N., Hingamp,P., Iudicone,D., Jaillon,O., Not,F., Ogata,H., Pesant,S., Speich,S., Stemmann,L., Sullivan,M.B., Weissenbach,J., Wincker,P., Karsenti,E., Raes,J., Acinas,S.G. and Bork,P. (2015) Structure and function of the global ocean microbiome. Science, 348(6237), 1261359.

Titus Brown,C. and Irber,L. (2016) sourmash: a library for MinHash sketching of DNA. The Journal of Open Source Software, 1(5), 27.

Truong,D.T., Franzosa,E.A., Tickle,T.L., Scholz,M., Weingart,G., Pasolli,E., Tett,A., Huttenhower,C. and Segata,N. (2015) MetaPhlAn2 for enhanced metagenomic taxonomic profiling. Nature Methods, 12(10), 902–903.

Viklund,J., Martijn,J., Ettema,T.J.G. and Andersson,S.G.E. (2013) Comparative and phylogenomic evidence that the alphaproteobacterium HIMB59 is not a member of the oceanic SAR11 clade. PLoS ONE, 8(11), e78858.

Wood,D.E. and Salzberg,S.L. (2014) Kraken: ultrafast metagenomic sequence classification using exact alignments. Genome biology, 15(3), R46.

Zhou,X., Shen,X., Hittinger,C.T. and Rokas,A. (2018) Evaluating Fast Maximum Likelihood-Based Phylogenetic Programs Using Empirical Phylogenomic Data Sets. Molecular Biology And Evolution, 35(2), 486–503.

