## Supplementary File for "PhyloMagnet: Fast and accurate screening of short-read meta-omics data using gene-centric phylogenetics"

**References** **18**

### Download and prepare Reference sequences

#### Prepare initial rp16 references

Download initial references from EggNOG and create multiple sequence alignments and trees. Save the reference packages in compressed form in `rp16_rpkg`.

```
singularity pull --name PhyloMagnet.simg shub://maxemil/PhyloMagnet:latest
singularity exec PhyloMagnet.simg python3 -c "import ete3; ncbi = ete3.NCBITaxa()"

nextflow run maxemil/PhyloMagnet \
            -with-singularity  PhyloMagnet.simg \
            --align_method 'mafft-einsi' \
            --phylo_method 'iqtree' \
            --cpus 36 \
            --megan_vmoptions "../MEGAN.vmoptions" \
            --reference_classes MBARC/eggnog_rp16.txt
            --reference_dir rp16_references
```

```
bash $HOME/.nextflow/assets/maxemil/PhyloMagnet/utils/make_reference_packages.sh \
            rp16_references rp16_rpkg
```

Create HMM models for the alignments of reference sequences to search additional genomes and complement the reference OGs.

```
mkdir rp16_hmms

for cog in rp16_references/C*/COG*[0-9].unique.aln;
do
  singularity exec -B $PWD:$PWD benchmark.simg hmmbuild rp16_hmms/$(basename
      ${cog%%.unique.aln}).hmm $cog;
done
```

#### Create the extended rp16 references including all genera included in the MBARC-26 dataset

Download the genomes of relatives of the MBARC species, annotate each sequence with the taxid

```
download(){
  while read id tax;
  do
    ass=$(wget -q
        "https://www.ncbi.nlm.nih.gov/entrez/eutils/esearch.fcgi?db=assembly&term="$id
        -O - | xml_grep -cond Id --text_only)
    ftp=$(wget -q
        "https://www.ncbi.nlm.nih.gov/entrez/eutils/esummary.fcgi?db=assembly&id="$ass
        -O - | xml_grep -cond FtpPath_GenBank --text_only)
    base=$(basename $ftp)
    wget -q $ftp/$base"_protein.faa.gz" -O $id.faa.gz
    wget -q $ftp/$base"_genomic.fna.gz" -O $id.fna.gz
    zcat "$f.faa.gz" | sed 's/>/>'"$tax"'\./g' | gzip > "$f.labelled.faa.gz" ;
  done < $1

  zcat *.labelled.faa.gz | gzip > all_proteomes.faa.gz
}

cd MBARC/genomes_relatives
download taxids.txt
cd ../..
```

Use the HMM to search for sequences in the relatives' genomes and add these sequences to the reference fasta

```
mkdir rp16_added_fasta

for hmm in rp16_hmms/COG*[0-9].hmm;
do
  singularity exec -B $PWD:$PWD benchmark.simg hmmsearch -T 50 --tblout
      rp16_added_fasta/$(basename ${hmm%%.hmm}).out $hmm
      MBARC/genomes_relatives/all_proteomes.faa.gz;
```

```
  grep -v "^#" rp16_added_fasta/$(basename ${hmm%%.hmm}).out | awk '{print $1}' |
      esl-sfetch -f MBARC/genomes_relatives/all_proteomes.faa.gz - >
      rp16_added_fasta/$(basename ${hmm%%.hmm}).hits.fasta ;
done

for cog in  rp16_references/C*/COG*[0-9].fasta;
do
  cat $cog rp16_added_fasta/$(basename ${cog%%.fasta}).hits.fasta >
      rp16_added_fasta/$(basename ${cog%%.fasta})_add.fasta
done
```

Use PhyloMagnet to create alignments and trees for each of the extended reference set. Package the files into reference packages (rpkg).

```
nextflow run maxemil/PhyloMagnet \
            -with-singularity PhyloMagnet.simg \
            --align_method 'mafft-einsi' \
            --phylo_method 'iqtree' \
            --cpus 36 \
            --megan_vmoptions "../MEGAN.vmoptions" \
            --local_ref "rp16_added_fasta/*_add.fasta" \
            --reference_dir rp16_added_references -resume

bash $HOME/.nextflow/assets/maxemil/PhyloMagnet/utils/make_reference_packages.sh
    rp16_added_references rp16_added_rpkg
```

Analogous to the rpkgs, create gpkgs to be used with graftM

```
mkdir rp16_added_gpkg

for cog in rp16_added_references/COG*;
do
  base=$(basename $cog)
  singularity exec -B $PWD:$PWD GraftM/graftM.img graftM create \
            --sequences $cog/$base.unique.fasta \
            --alignment $cog/$base.unique.aln \
            --rerooted_tree $cog/$base.treefile \
            --taxonomy $cog/$base.taxid.map \
            --output rp16_added_gpkg/$base.gpkg
done
```

#### Get and prepare Chloroplast gene references from uniprot.

Print the FASTA header in the format TAXID.ACCESSION, similar to how EggNOG references are formatted.

```
clean_headers(){
  python3 <<<"""
from Bio import SeqIO
import re

re.compile('OX=[0-9]*')
pattern = re.compile('OX=[0-9]*')
```

```

for gene in ['atpA', 'atpB', 'petB', 'petD', 'psaA', 'psaB', 'psbA', 'psbB', 'psbC', 'psbD', 'psbE', 'psbI']:
    recs = []
    for rec in SeqIO.parse('{}.fasta'.format(gene),'fasta'):
        seqid = rec.id.split('|')[1]
        seqtax = pattern.search(rec.description).group(0).split('=')[1]
        rec.id = '{}.{}'.format(seqtax, seqid)
        rec.description = ''
        recs.append(rec)
    with open('{}.fasta'.format(gene), 'w') as outhandle:
        SeqIO.write(recs, outhandle, 'fasta')
  """
}

mkdir chloroplast_references_uniprot
cd chloroplast_references_uniprot

for gene in "atpA" "atpB" "petB" "petD" "psaA" "psaB" "psbA" "psbB" "psbC" "psbD" "psbE" "psbI";
do
  wget "https://www.uniprot.org/uniprot/?query=gene%3A\
  $gene+(reviewed%3Ayes+OR+dinophyceae)+(chloroplast+OR+plastid)&format=fasta" \
-O "$gene.fasta"
done
clean_headers
cd ..
```

Reconstruct alignmnents and trees for chloroplast genes and package them into rpkgs

```
nextflow run maxemil/PhyloMagnet \
            -with-singularity PhyloMagnet.simg \
            --cpus 40 \
            --local_ref "chloroplast_references_uniprot/*.fasta" \
            --megan_vmoptions "../MEGAN.vmoptions" \
            --phylo_method 'iqtree' \
            --align_method 'mafft-einsi' \
            --reference_dir 'chloroplast_references'

bash $HOME/.nextflow/assets/maxemil/PhyloMagnet/utils/make_reference_packages.sh \
            chloroplast_references/ chloroplast_rpkgs/
```

### Download and prepare query datasets

Python script to download sra files from ENA and subsequently convert them to FASTQ files (basically taken from the template used in PhyloMagnet)

```
donwload_fastq(){
  python3 <<<"""
import requests
import shutil
```

```
import subprocess

acc = '$1'
url = ''
if len(acc) == 9:
    url = 'http://ftp.sra.ebi.ac.uk/vol1/{}/{}/{}'.format(acc[0:3].lower(), acc[0:6],
        acc)
elif len(acc) == 10:
    url = 'http://ftp.sra.ebi.ac.uk/vol1/{}/{}/{}/{}'.format(acc[0:3].lower(), acc[0:6],
        "00" + acc[-1], acc)
elif len(acc) == 11:
    url = 'http://ftp.sra.ebi.ac.uk/vol1/{}/{}/{}/{}'.format(acc[0:3].lower(), acc[0:6],
        "0" + acc[-2:], acc)

r = requests.get(url, stream=True)
with open('$1.sra', 'wb') as f:
      shutil.copyfileobj(r.raw, f)
  """
  fastq-dump --gzip --readids --split-spot --skip-technical --clip $1.sra
}
```

#### MBARC-26

Download MBARC-26 Illumina metagenomic dataset from ENA

```
mkdir MBARC/fastq
cd MBARC/fastq
donwnload_fastq SRR3656745
```

Subsample 1% and 10% from the MBARC-26 data for benchmarking

```
seqtk sample -s11 <(gunzip -c SRR3656745.fastq.gz) 0.01 | pigz -9 >
    SRR3656745.1perc.fastq.gz
seqtk sample -s11 <(gunzip -c SRR3656745.fastq.gz) 0.1 | pigz -9 >
    SRR3656745.10perc.fastq.gz
```

#### Tara Oceans

Download Tara Oceans metagenomic data from ENA

```
mkdir Tara_Southern_Ocean/fastq
cd Tara_Southern_Ocean/fastq

for id in ERR598945 ERR599008 ERR599059 ERR599090 ERR599104 ERR599121 ERR599125
    ERR599176;do
  donwnload_fastq $id
done
```

##### Coral Bleaching

Download Coral Bleaching metatranscriptomic data from ENA, concatenate them to a single dataset.

```
mkdir Tara_Southern_Ocean/fastq
cd Tara_Southern_Ocean/fastq

for id in SRR5453739 SRR5453740 SRR5453741 SRR5453742 SRR5453743 SRR5453744 SRR5453745
    SRR5453746 SRR5453747 SRR5453748 SRR5453749 SRR5453750 SRR5453751 SRR5453752
    SRR5453753 SRR5453754 SRR5453755 SRR5453756 SRR5453757 SRR5453758 SRR5453759
    SRR5453760 SRR5453761 SRR5453762 SRR5453763 SRR5453764 SRR5453765;do
  donwnload_fastq $id
done
cat SRR*.fastq.gz > PRJNA377366.fastq.gz
```

#### Download published results as comparison for benchmarking

##### MBARC-26

Compare PhyloMagnet results to GraftM results, see file `MBARC/genomes_mapped.txt` for genome mapping information (i.e. abundance).

##### Tara Oceans

```
cd Tara_Southern_Ocean
wget https://ndownloader.figshare.com/files/8243654 -O NON_REDUNDANT_MAGs.tar.gz
tar -xzf NON_REDUNDANT_MAGs.tar.gz
# similarly, download raw BINs
```

Compute taxonomic labels for additional genome bins using sourmash (download the database from sourmash's website https://sourmash.readthedocs.io/en/latest/databases.html)q

```
cd sourmash
wget https://osf.io/nemkw/download -O t-genbank-k51.lca.json.gz
nextflow run sourmash.nf --genomes ../NON_REDUNDANT_MAGs/* --reference
    genbank-k51.lca.json.gz --outdir signatures-MAGS -qs 35

cd ..
```

##### Coral Bleaching

```
mkdir Coral_Bleaching/transcriptome
cd Coral_Bleaching/transcriptome

split_alignment(){
  python3 <<<"""
from Bio import SeqIO
refs = [rec.id for rec in SeqIO.parse('$2$1.unique.aln', 'fasta')]
queries = [rec for rec in SeqIO.parse('$1.refquer.aln', 'fasta') if not rec.id in refs]
```

```
with open('$1.quer.aln', 'w') as out:
    for rec in queries:
        SeqIO.write(rec, out, 'fasta')
"""
}
```

Download the metatranscriptomic assembly from GEO database. prepare blast database

```
TRANSCRIPTOME_FILE=GSE97888%5FMetatranscriptome%5Fqualityfiltered%2Efasta%2Egz
wget
    https://www.ncbi.nlm.nih.gov/geo/download/?acc=GSE97888&format=file&file=$TRANSCRIPTOME_FILE
    -O GSE97888.fasta.gz
gunzip GSE97888.fasta.gz
makeblastdb -in GSE97888.fasta -out GSE97888.db -dbtype nucl
```

For each reference OG, identify homologous transcripts and place them onto the ref tree.

```
cd ..
mkdir place_transcripts
cd place_transcripts

for dir in ../references/*/;
do
  gene=${dir#../references/}
  gene=${gene%/}
  tblastn -db ../transcriptome/GSE97888.db -num_threads 30 -query $dir$gene.unique.fasta
      -evalue 1e-15 -outfmt '6 qseqid sseqid pident length mismatch gapopen qstart qend
      sstart send sframe evalue bitscore' -out $gene.tblastn
  python3 ../get_blast_hsp.py -b $gene.tblastn -f ../transcriptome/GSE97888.fasta -o
      $gene.hits --translate -n qseqid sseqid pident length mismatch gapopen qstart qend
      sstart send sframe evalue bitscore
  python3 ../get_blast_hsp.py -b $gene.tblastn -f ../transcriptome/GSE97888.fasta -o
      $gene.nucl.hits -n qseqid sseqid pident length mismatch gapopen qstart qend sstart
      send sframe evalue bitscore
  singularity exec -B /local:/local ../../PhyloMagnet.simg trimal -in
      $dir$gene.unique.aln -out $gene.ref.phy -phylip
  singularity exec -B /local:/local ../../PhyloMagnet.simg papara -t $dir$gene.treefile
      -s $gene.ref.phy -q $gene.hits -a -n $gene -r
  singularity exec -B /local:/local ../../PhyloMagnet.simg epa-ng --split $gene.ref.phy
      papara_alignment.$gene
  mv query.fasta $gene.quer.aln
  singularity exec -B /local:/local ../../PhyloMagnet.simg epa-ng --ref-msa
      $dir$gene.unique.aln --tree $dir$gene.treefile --query $gene.quer.aln --model
      $dir$gene.modelfile --no-heur --threads 10
  mv epa_result.jplace $gene.jplace
  mv epa_info.log $gene.epa_info.log
  singularity exec -B /local:/local ../../PhyloMagnet.simg gappa analyze assign
      --jplace-path $gene.jplace --taxon-file $dir$gene.taxid.map --threads 10
  mv profile.csv $gene.csv
  rm labelled_tree
  mv per_pquery_assign $gene.assign
  singularity exec -B /local:/local ../../PhyloMagnet.simg gappa analyze graft
      --name-prefix 'Q_' --jplace-path $gene.jplace --threads 10
```

```
done
cd ../..
```

### Run Benchmarks

#### MBARC-26 benchmark

Run and time PhyloMagnet on MBARC data (full, 10% and 1% subsampled)

```
/usr/bin/time --output=queries_1perc_timed.txt -v nextflow run maxemil/PhyloMagnet \
            --lineage "genus" \
            -with-singularity  ../PhyloMagnet.simg \
            --cpus 10 \
            --reference_packages "../rp16_added_rpkg/*.tgz*" \
            --fastq "fastq/SRR3656745.1perc.fastq.gz" \
            --megan_vmoptions "../MEGAN.vmoptions" \
            -with-report MBARC_report_1perc.html \
            --queries_dir queries_1perc

/usr/bin/time --output=queries_10perc_timed.txt -v nextflow run maxemil/PhyloMagnet \
            --lineage "genus" \
            -with-singularity  ../PhyloMagnet.simg \
            --cpus 10 \
            --reference_packages "../rp16_added_rpkg/*.tgz*" \
            --fastq "fastq/SRR3656745.10perc.fastq.gz" \
            --megan_vmoptions "../MEGAN.vmoptions" \
            -with-report MBARC_report_10perc.html \
            --queries_dir queries_10perc

/usr/bin/time --output=queries_timed.txt -v nextflow run maxemil/PhyloMagnet \
            --lineage "genus" \
            -with-singularity  ../PhyloMagnet.simg \
            --cpus 10 \
            --reference_packages "../rp16_added_rpkg/*.tgz*" \
            --fastq "fastq/SRR3656745.fastq.gz" \
            --megan_vmoptions "../MEGAN.vmoptions" \
            -with-report MBARC_report.html \
            --queries_dir queries

# --lineage "Clostridium","Ruminiclostridium","Coraliomargarita",\
#           "Corynebacterium","Desulfosporosinus","Desulfotomaculum",\
#           "Echinicola","Escherichia","Fervidobacterium","Frateuria",\
#           "Halovivax","Hirschia","Meiothermus","Natronobacterium",\
#           "Natronococcus","Nocardiopsis","Olsenella","Pseudomonas",\
#           "Salmonella","Segniliparus","Sediminispirochaeta","Streptococcus",\
#           "Terriglobus","Thermobacillus","Enterobacteriaceae" \
```

Run and time GraftM on MBARC data (full, 10% and 1% subsampled)

```
mkdir GraftM_output_1perc
```

```
/usr/bin/time --output=graftM_1perc_timed.txt -v bash MBARC_GraftM.sh
    "MBARC/fastq/SRR3656745.1perc.fastq.gz" "GraftM_output_1perc"
mkdir GraftM_output_10perc
/usr/bin/time --output=graftM_10perc_timed.txt -v bash MBARC_GraftM.sh
    "MBARC/fastq/SRR3656745.10perc.fastq.gz" "GraftM_output_10perc"
mkdir GraftM_output_timed
/usr/bin/time --output=graftM_timed.txt -v bash MBARC_GraftM.sh
    "MBARC/fastq/SRR3656745.fastq.gz" "GraftM_output_timed"
```

Where `MBARC_GraftM.sh` look like this:

```
for gpkg in rp16_added_gpkg/*
do
  base=$(basename ${gpkg%%.gpkg})
  singularity exec -B $PWD:$PWD graftM.simg graftM graft \
                                              --forward $1 \
                                              --graftm_package $gpkg \
                                              --input_sequence_type nucleotide \
                                              --search_method hmmsearch \
                                              --assignment_method pplacer \
                                              --threads 10 \
                                              --verbosity 5 \
                                              --log $2/"$base"_GraftM.log \
                                              --output_directory $2/"$base"
done
```

#### Tara Southern Oceans Benchmark

Run PhyloMagnet on the Coral Bleaching metatranscriptomic dataset

```
nextflow run maxemil/PhyloMagnet \
            --lineage "family" \
            -with-singularity  ../PhyloMagnet.simg \
            --cpus 32 \
            --fastq "fastq/*.fastq.gz" \
            --reference_packages "../rp16_rpkg/*" \
            --megan_vmoptions "../MEGAN.vmoptions"
```

#### Coral Bleaching Benchmark

Run PhyloMagnet on the Tara Oceans metagenomic dataset

```
nextflow run maxemil/PhyloMagnet \
            --lineage "family","Dinophyceae" \
            -with-singularity  ../PhyloMagnet.simg \
            --cpus 40 \
            --reference_packages "../chloroplast_rpkgs/*" \
            --megan_vmoptions "../MEGAN.vmoptions" \
            --fastq "fastq/PRJNA377366.fastq.gz" \
            -with-report "Coral_Bleaching.html"
```

### Evaluate Benchmarks

#### Fig 2:

```
import matplotlib
matplotlib.use('Agg')
import pandas as pd
import seaborn as sns
from itertools import product
from collections import defaultdict
import matplotlib.pyplot as plt
import glob
from ete3 import ncbi_taxonomy

MBARC_genera = ['Clostridium','Ruminiclostridium','Coraliomargarita',
    'Corynebacterium','Desulfosporosinus','Desulfotomaculum','Echinicola',
    'Escherichia','Fervidobacterium','Frateuria','Halovivax','Hirschia',
    'Olsenella','Pseudomonas','Salmonella','Segniliparus','Sediminispirochaeta',
    'Meiothermus','Natronobacterium','Natronococcus','Nocardiopsis',
    'Streptococcus','Terriglobus','Thermobacillus']

def get_counts_tree_taxon(df):
    taxa = set(df["Taxon"])
    samples = set(df["Tree"])
    counts = pd.DataFrame(columns=samples, index=taxa)
    counts = counts.fillna(0)
    for t,s in product(taxa, samples):
        counts[s].loc[t] = df[(df["Tree"] == s) & (df["Taxon"] == t)].shape[0]
    return counts

def parse_graftM_results(path):
    ncbi = ncbi_taxonomy.NCBITaxa()
    lineages = defaultdict(lambda: defaultdict(int))
    rank = "genus"
    for infile in glob.glob("{}/COG*/combined_count_table.txt".format(path)):
        taxids = set()
        tree = infile.split('/')[1]
        for line in open(infile):
            lineage = [l.strip() for l in line.split('\t')[2].split(';')]
            name2tax = ncbi.get_name_translator(lineage)
            taxids |= set([taxid for l in name2tax.values() for taxid in l])
        for k,v in ncbi.get_rank(taxids).items():
                if v == rank:
                    lineages[tree][ncbi.get_taxid_translator([k])[k]] += 1
    return pd.DataFrame.from_dict(lineages).fillna(0)

def parse_PhyloMagnet_results(infile):
    df = pd.read_csv(infile, sep='\t', header=None, names=['Sample', 'Tree', 'Taxon',
        'value'], dtype=str)
    df = df[df['value'] == 'True']
    return get_counts_tree_taxon(df)
```

```python

def divide_tp_and_fp(df, tool):
    df = df.apply(lambda x: x if x.name in MBARC_genera else (-1 * x), axis=1)
    df_tp = pd.DataFrame({'counts':df[df >= 0].sum(), 'tool':tool})
    df_fp = pd.DataFrame({'counts':df[df <= 0].sum(), 'tool':tool})
    return df_tp.append(df_fp)

def lighten_color(color, amount=0.5):
    import matplotlib.colors as mc
    import colorsys
    try:
        c = mc.cnames[color]
    except:
        c = color
    c = colorsys.rgb_to_hls(*mc.to_rgb(c))
    return colorsys.hls_to_rgb(c[0], 1 - amount * (1 - c[1]), c[2])

def get_counts_both_tools():
    graftm_1 = parse_graftM_results("GraftM_output_1perc")
    df = divide_tp_and_fp(graftm_1, 'GraftM 1%')
    graftm_10 = parse_graftM_results("GraftM_output_10perc")
    df = df.append(divide_tp_and_fp(graftm_10, 'GraftM 10%'))
    graftm = parse_graftM_results("GraftM_output")
    df = df.append(divide_tp_and_fp(graftm, 'GraftM'))

    counts_1 = parse_PhyloMagnet_results("MBARC/queries_1perc/tree_decisions.txt")
    df = df.append(divide_tp_and_fp(counts_1, 'PhyloMagnet 1%'))
    counts_10 = parse_PhyloMagnet_results("MBARC/queries_10perc/tree_decisions.txt")
    df = df.append(divide_tp_and_fp(counts_10, 'PhyloMagnet 10%'))
    counts = parse_PhyloMagnet_results("MBARC/queries/tree_decisions.txt")
    df = df.append(divide_tp_and_fp(counts, 'PhyloMagnet'))
    return df

def plot_counts_fp_tp(df):
    clr_palette = [lighten_color(sns.xkcd_rgb['scarlet'], 0.7),
                   sns.xkcd_rgb['scarlet'],
                   lighten_color(sns.xkcd_rgb['scarlet'], 1.3),
                   lighten_color(sns.xkcd_rgb['denim'], 0.7),
                   sns.xkcd_rgb['denim'],
                   lighten_color(sns.xkcd_rgb['denim'], 1.3)]

    fig, ax = plt.subplots(figsize=(10,6), tight_layout=True)
    sns.swarmplot(x=df.index, y='counts', data=df, hue='tool',
                   ax=ax, palette=clr_palette, linewidth=0.4, size=6)

    ax.set_xticklabels(labels=df.index, rotation=90)
    ax.set_ylabel('True and False positive')
    ax.set_ylim(-16, 25)
    ax.axhline(0, color=sns.xkcd_rgb['denim'])
    ax.legend(frameon=False, bbox_to_anchor=(1, 1), loc=2)
    fig.savefig('Fig2.pdf', orientation='landscape', dpi=500)
```

```

if __name__ == '__main__':
    df = get_counts_both_tools()
    plot_counts_fp_tp(df)
```

#### Fig 3:

This figure is simply the heatmap taken from the ouput of the Tara oceans benchmark, added with a ‘taxonomic tree’ as it can be found on ncbi. `Tara_Southern_Ocean/queries/decision_heatmap.pdf`

#### Fig 4:

These are the two trees `Coral_Bleaching/queries/PRJNA377366/PRJNA377366-psbb.newick` and `Coral_Bleaching/place_transcripts/psbb.newick` aligned to each other so as to be able to compare the placements

#### Fig S1:

```
import matplotlib
matplotlib.use('Agg')
import pandas as pd
import seaborn as sns
from itertools import product
from collections import defaultdict
import matplotlib.pyplot as plt
import glob
from ete3 import ncbi_taxonomy

def get_counts_sample_taxon(df):
    taxa = set(df["Taxon"])
    samples = set(df["Sample"])
    counts = pd.DataFrame(columns=samples, index=taxa)
    counts = counts.fillna(0)
    for t,s in product(taxa, samples):
        counts[s].loc[t] = df[(df["Sample"] == s) & (df["Taxon"] == t)].shape[0]
    return counts

def parse_graftM_results_sample(path):
    ncbi = ncbi_taxonomy.NCBITaxa()
    lineages = defaultdict(int)
    rank = "genus"
    for infile in glob.glob("{}/COG*/combined_count_table.txt".format(path)):
        taxids = set()
        for line in open(infile):
            lineage = [l.strip() for l in line.split('\t')[2].split(';')]
            name2tax = ncbi.get_name_translator(lineage)
            taxids |= set([taxid for l in name2tax.values() for taxid in l])
        for k,v in ncbi.get_rank(taxids).items():
                if v == rank:
                    lineages[ncbi.get_taxid_translator([k])[k]] += 1
```

```
    return pd.DataFrame.from_dict(lineages,orient='index')

def parse_PhyloMagnet_results_sample(infile):
    df = pd.read_csv(infile, sep='\t', header=None, names=['Sample', 'Tree', 'Taxon',
        'value'], dtype=str)
    df = df[df['value'] == 'True']
    return get_counts_sample_taxon(df)

def get_counts_both_tools_sample():
    phylomagnet = parse_PhyloMagnet_results_sample("MBARC/queries/tree_decisions.txt")
    graftm = parse_graftM_results_sample("GraftM_output")
    return (phylomagnet, graftm)

def plot_heatmap(phylomagnet, graftm):
    fig, ax = plt.subplots(figsize=(10,20), tight_layout=True)
    compare = pd.DataFrame({'GraftM':graftm[0],'PhyloMagnet':phylomagnet.SRR3656745})
    compare = compare.fillna(0)
    compare = compare.sort_values(by=['PhyloMagnet', 'GraftM'],ascending=False)
    sns.heatmap(compare, annot=True, cmap='Reds', xticklabels=True, ax=ax)
    fig.savefig('FigS1.pdf')

if __name__ == '__main__':
    phylomagnet, graftm = get_counts_both_tools_sample()
    plot_heatmap(phylomagnet, graftm)
```

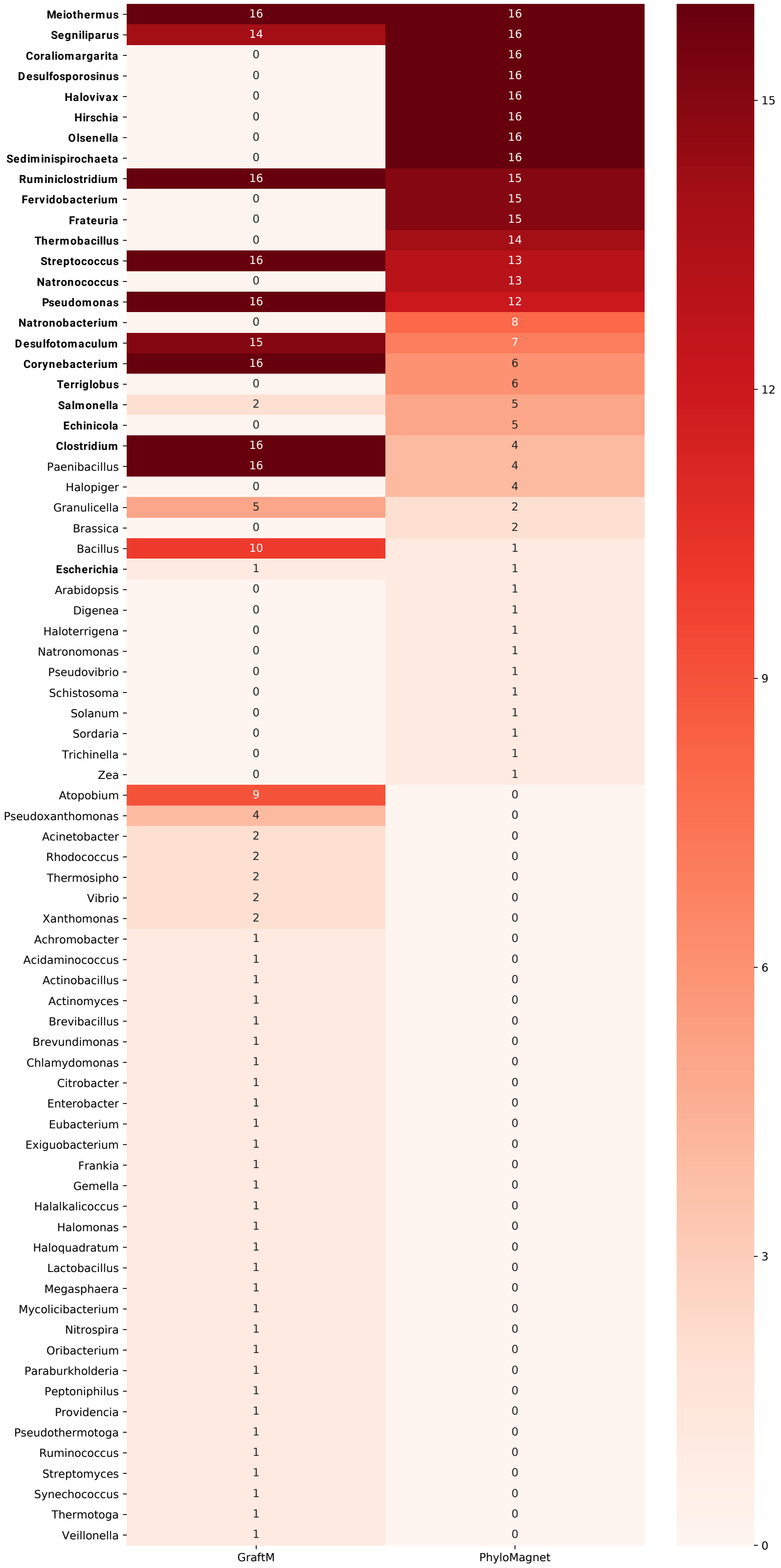

Figure S1: Classification results of PhyloMagnet and GraftM on the full MBARC-26 dataset. Values and colors correspond to the number of rp16 trees a genus was identified in. Those genera that are part of the MBARC-26 community are written in bold.

#### Fig S2:

```python
import matplotlib.pyplot as plt
import pandas as pd
import seaborn as sns

fig, ax = plt.subplots(2, 1, figsize=(20,10), tight_layout=True)

df = pd.DataFrame.from_csv('MBARC/runtimes.csv',index_col=None)
sns.catplot(x='size', y='time',hue='tool', data=df, ax=ax[0], kind="bar",
    palette="muted", legend=False)
ax[0].set(xlabel="size [Gb]", ylabel='time [s]')

df = pd.DataFrame.from_csv('MBARC/memory.csv',index_col=None)
sns.catplot(x='size', y='memory',hue='tool', data=df, ax=ax[1], kind="bar",
    palette="muted", legend_out=True)
ax[1].set(xlabel="size [Gb]", ylabel='memory [GB]')

fig.savefig('FigS2.pdf')
```

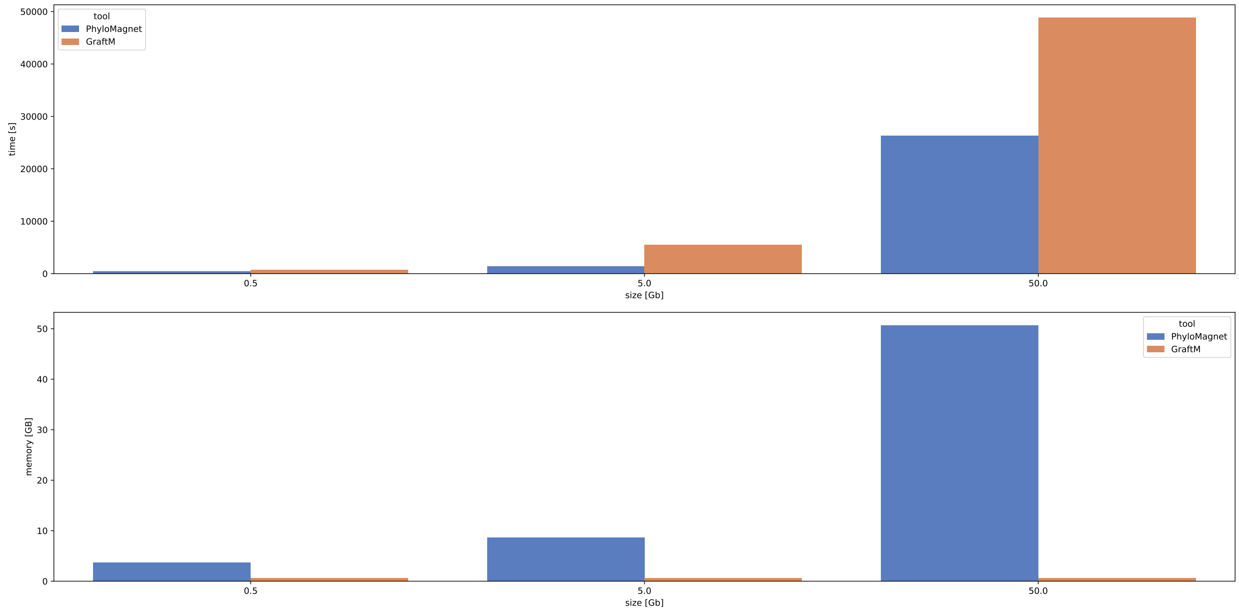

Figure S2: Computational footprint of PhyloMagnet and GraftM for the complete MBARC-26 dataset (50Gb), the subsample of 1% (5Gb) and of 10% (0.5Gb). top: runtime on 10 CPUs. bottom: peak memory usage.

#### Table S1:

Table S1: Organisms included in the MBARC-26 dataset. For each organism, current taxonomy (species, genus and family as given in the ncbi taxonomy), ncbi assembly ID and percentage mapped illumina reads as presented in Singer *et al.* (2018) are shown.

| genus | species | family | Assembly | TaxID | mapped reads [%] |
|---|---|---|---|---|---|
| Desulfosporosinus | acidiphilus | Peptococcaceae | NC_018068 | 646529 | 15.11 |
| Sediminispirochaeta | smaragdinae | Spirochaetaceae | NC_014364 | 573413 | 11.39 |
| Fervidobacterium | pennivorans | Fervidobacteriaceae | NC_017095 | 771875 | 11.26 |
| Meiothermus | Silvanus | Thermaceae | NC_014212 | 526227 | 8.56 |
| Thermobacillus | composti | Paenibacillaceae | NC_019897 | 717605 | 8.5 |
| Hirschia | baltica | Hyphomonadaceae | NC_012982 | 582402 | 8.16 |
| Desulfotomaculum | gibsoniae | Peptococcaceae | NC_021184 | 767817 | 6.91 |
| Desulfosporosinus | meridiei | Peptococcaceae | NC_018515 | 768704 | 4.61 |
| Frateuria | aurantia | Rhodanobacteraceae | NC_017033 | 767434 | 3.99 |
| Natronococcus | occultus | Natrialbaceae | NC_019974.1 | 694430 | 3.55 |
| Coraliomargarita | akajimensis | Puniceicoccaceae | NC_014008 | 583355 | 3.41 |
| Natronobacterium | gregoryi | Natrialbaceae | NC_019792.1 | 797304 | 2.46 |
| Olsenella | uli | Atopobiaceae | NC_014363 | 633147 | 2.26 |
| Terriglobus | roseus | Acidobacteriaceae | NC_018014 | 926566 | 2.07 |
| Halovivax | ruber | Natrialbaceae | CP003050.1 | 797302 | 1.75 |
| Pseudomonas | stutzeri | Pseudomonadaceae | NC_019936 | 644801 | 1.55 |
| Segniliparus | rotundus | Segniliparaceae | NC_014168 | 640132 | 1.41 |
| Echinicola | vietnamensis | Cyclobacteriaceae | NC_019904 | 926556 | 0.62 |
| Salmonella | enterica | Enterobacteriaceae | NC_010067 | 41514 | 0.52 |
| Ruminiclostridium | thermocellum | Ruminococcaceae | NC_009012 | 203119 | 0.43 |
| Streptococcus | pyogenes | Streptococcaceae | NC_002737 | 160490 | 0.43 |
| Clostridium | perfringens | Clostridiaceae | NC_008261 | 195103 | 0.42 |
| Corynebacterium | glutamicum | Corynebacteriaceae | NC_003450 | 196627 | 0.3 |
| Escherichia | coli | Enterobacteriaceae | NC_000913 | 511145 | 0.18 |
| Salmonella | bongori | Enterobacteriaceae | NC_015761 | 218493 | 0.14 |
| Nocardiopsis | dassonvillei | Nocardiopsaceae | NC_014211 | 446468 | 0 |

#### Table S2:

Table S2: Taxonomic annotation of MAGs extracted from the Tara southern oceans dataset by Delmont *et. al* (2018). The inferred taxonomic labels from domain to species level (as far as available) are provided for the 11 prokaryotic nonredundant MAGs (inferred with GraftM; annotation of TARA_ANE_MAG_00007 is given for TARA_SOC_MAG_00009, as they have ANI >99%) as well as for those additional raw bins where a taxonomic annotation could be inferred with sourmash.

| MAG/Bin | Phylum | Class | Order | Family |
|---|---|---|---|---|
| TARA_SOC_MAG_00001 | Proteobacteria | Alphaproteobacteria | - | - |
| TARA_SOC_MAG_00002 | Proteobacteria | Gammaproteobacteria | Alteromonadales | Alteromonadaceae |
| TARA_SOC_MAG_00003 | Bacteroidetes | Flavobacteriia | Flavobacteriales | - |
| TARA_SOC_MAG_00004 | Bacteroidetes | Flavobacteriia | Flavobacteriales | - |
| TARA_SOC_MAG_00005 | Bacteroidetes | Flavobacteriia | Flavobacteriales | - |
| TARA_SOC_MAG_00006 | Bacteroidetes | Flavobacteriia | Flavobacteriales | **Cryomorphaceae** |
| TARA_SOC_MAG_00007 | Proteobacteria | Gammaproteobacteria | - | - |
| TARA_SOC_MAG_00008 | Proteobacteria | Alphaproteobacteria | Rickettsiales | - |
| TARA_SOC_MAG_00009 | Actinobacteria | Actinobacteria | Actinomycetales | Microbacteriaceae |
| TARA_SOC_MAG_00010 | Proteobacteria | Gammaproteobacteria | - | - |
| TARA_SOC_MAG_00011 | Bacteroidetes | Flavobacteriia | Flavobacteriales | Flavobacteriaceae |
| TARA_SOC_MAG_00012 | Bacteroidetes | Flavobacteriia | Flavobacteriales | - |
| TARA_SOC_Bin_00046 | Bacteroidetes | Flavobacteriia | Flavobacteriales | Flavobacteriaceae |
| TARA_SOC_Bin_00048 | Proteobacteria | Alphaproteobacteria | Pelagibacterales | Pelagibacteraceae |
| TARA_SOC_Bin_00060 | Bacteroidetes | Flavobacteriia | Flavobacteriales | Flavobacteriaceae |
| TARA_SOC_Bin_00063 | Proteobacteria | Gammaproteobacteria | Cellvibrionales | Porticoccaceae |
| TARA_SOC_Bin_00074 | Proteobacteria | Alphaproteobacteria | Pelagibacterales | Pelagibacteraceae |
| TARA_SOC_Bin_00075 | Proteobacteria | Gammaproteobacteria | Cellvibrionales | Porticoccaceae |
| TARA_SOC_Bin_00095 | Bacteroidetes | Flavobacteriia | Flavobacteriales | Cryomorphaceae |
| TARA_SOC_Bin_00115 | Proteobacteria | Gammaproteobacteria | Cellvibrionales | Porticoccaceae |
| TARA_SOC_Bin_00226 | Proteobacteria | Gammaproteobacteria | Cellvibrionales | Porticoccaceae |
| TARA_SOC_Bin_00242 | Proteobacteria | Alphaproteobacteria | Rhodobacterales | Rhodobacteraceae |

Table S2: continued

| MAG/Bin | Genus | Species | Tool |
|---|---|---|---|
| TARA_SOC_MAG_00001 | - | - | GraftM |
| TARA_SOC_MAG_00002 | - | - | GraftM |
| TARA_SOC_MAG_00003 | - | - | GraftM |
| TARA_SOC_MAG_00004 | - | - | GraftM |
| TARA_SOC_MAG_00005 | - | - | GraftM |
| TARA_SOC_MAG_00006 | **unassigned** | **Cryomorphaceae bacterium ASP10-05a** | GraftM/**sourmash** |
| TARA_SOC_MAG_00007 | - | - | GraftM |
| TARA_SOC_MAG_00008 | - | - | GraftM |
| TARA_SOC_MAG_00009 | Microbacterium | - | GraftM |
| TARA_SOC_MAG_00010 | - | - | GraftM |
| TARA_SOC_MAG_00011 | Polaribacter | - | GraftM |
| TARA_SOC_MAG_00012 | - | - | GraftM |
| TARA_SOC_Bin_00046 | - | Flavobacteriaceae bacterium ASP10-09a | sourmash |
| TARA_SOC_Bin_00048 | Candidatus Pelagibacter | Candidatus Pelagibacter sp. IMCC9063 | sourmash |
| TARA_SOC_Bin_00060 | Flavobacterium | Flavobacterium sp. SCGC AAA160-P02 | sourmash |
| TARA_SOC_Bin_00063 | - | marine gamma proteobacterium ASP10-03a | sourmash |
| TARA_SOC_Bin_00074 | Candidatus Pelagibacter | Candidatus Pelagibacter sp. IMCC9063 | sourmash |
| TARA_SOC_Bin_00075 | - | marine gamma proteobacterium ASP10-03a | sourmash |
| TARA_SOC_Bin_00095 | - | Cryomorphaceae bacterium ASP10-05a | sourmash |
| TARA_SOC_Bin_00115 | - | marine gamma proteobacterium ASP10-03a | sourmash |
| TARA_SOC_Bin_00226 | - | marine gamma proteobacterium ASP10-03a | sourmash |
| TARA_SOC_Bin_00242 | - | Rhodobacteraceae bacterium ASP10-04a | sourmash |

### References

Altschul, S. F., Madden, T. L., Schäffer, A. A., Zhang, J., Zhang, Z., Miller, W., & Lipman, D. J. (1997). Gapped BLAST and PSI-BLAST: a new generation of protein database search programs. Nucleic Acids Research, 25(17), 3389–402. Retrieved from http://www.ncbi.nlm.nih.gov/pubmed/9254694

Boyd, J. A., Woodcroft, B. J., & Tyson, G. W. (2018). GraftM: a tool for scalable, phylogenetically informed classification of genes within metagenomes. Nucleic Acids Research, 46(10), e59–e59. https://doi.org/10.1093/nar/gky174

Di Tommaso, P., Chatzou, M., Floden, E. W., Barja, P. P., Palumbo, E., & Notredame, C. (2017). Nextflow enables reproducible computational workflows. Nature Biotechnology, 35(4), 316–319. https://doi.org/10.1038/nbt.3820

Eddy, S. R. (2011). Accelerated Profile HMM Searches. PLoS Computational Biology, 7(10), e1002195. https://doi.org/10.1371/journal.pcbi.1002195

Katoh, K., & Standley, D. M. (2013). MAFFT multiple sequence alignment software version 7: Improvements in performance and usability. Molecular Biology and Evolution, 30(4), 772–780. https://doi.org/10.1093/molbev/mst010

Nguyen, L. T., Schmidt, H. A., Von Haeseler, A., & Minh, B. Q. (2015). IQ-TREE: A fast and effective stochastic algorithm for estimating maximum-likelihood phylogenies. Molecular Biology and Evolution, 32(1), 268–274. https://doi.org/10.1093/molbev/msu300

Titus Brown, C., & Irber, L. (2016). sourmash: a library for MinHash sketching of DNA. The Journal of Open Source Software, 1(5), 27. https://doi.org/10.21105/joss.00027
